# Gene Therapy Reforms Photoreceptor Structure and Restores Vision in *NPHP5*-associated Leber Congenital Amaurosis

**DOI:** 10.1101/2020.10.07.329821

**Authors:** Gustavo D. Aguirre, Artur V. Cideciyan, Valérie L. Dufour, Ana Ripolles García, Raghavi Sudharsan, Malgorzata Swider, Roman Nikonov, Simone Iwabe, Sanford L. Boye, William W. Hauswirth, Samuel G. Jacobson, William A. Beltran

## Abstract

The inherited childhood blindness caused by mutations in *NPHP5*, a form of Leber congenital amaurosis, results in abnormal development, dysfunction and degeneration of photoreceptors. A naturally occurring *NPHP5* mutation in dogs results in a phenotype that very nearly duplicates the human retinopathy in terms of the photoreceptors involved, spatial distribution of degeneration and the natural history of vision loss. We show that AAV-mediated *NPHP5* gene augmentation of mutant canine retinas at the time of active degeneration and peak cell death stably restores photoreceptor structure, function, and vision with either the canine or human *NPHP5* transgenes. Mutant cone photoreceptors, which failed to form outer segments during development, reform this structure after treatment. Degenerating rod photoreceptor outer segments are stabilized and develop normal structure. This process begins within 8 weeks following treatment, and remains stable throughout the 6 month post treatment period. In both photoreceptor cell classes, mislocalization of rod and cone opsins is minimized or reversed. Retinal function and functional vision are restored. Efficacy of gene therapy in this large animal ciliopathy model of Leber congenital amaurosis provides a path for translation to human treatment.

## Introduction

The hereditary childhood blindness, Leber congenital amaurosis (LCA), is a heterogeneous group of disorders affecting the photoreceptor cells or retinal pigment epithelium (RPE) that are characterized by severe visual impairment or early-onset childhood blindness, and currently considered to be mostly untreatable (*1*). To date, disease causing gene mutations have been identified in autosomal recessive (n=23) or dominant (n=3) forms of LCA (https://sph.uth.edu/retnet/sum-dis.htm#B-diseases; September 1, 2020). These genes play essential but varied roles in outer retinal function and vision, ranging from vitamin A isomerization in the visual cycle, phototransduction, photoreceptor development and maintenance, and intracellular transport, among others ((*2*) and see(*3*) for review). Of these, mutations in genes involved in transport through the photoreceptor ciliary transition zone (aka connecting cilium) result in retinal ciliopathies, and comprise a major subclass of LCA with at least 8 genes (https://sph.uth.edu/retnet/sum-dis.htm#B-diseases (*3*–*6*)) as well as members of the BBSome complex (*7*–*9*). In addition to the retinal phenotype, CNS abnormalities and/or renal cystic disease can occur. One such ciliopathy resulting from *NPHP5* (*IQCB1*) mutations, Senior Loken Syndrome (SLSN), includes nephronophthisis (NPHP) and LCA with the cystic kidney disease being the most frequent cause of end-stage renal failure in the first three decades of life (*10*).

*NPHP5* mutations represent a rare form of retinal disease estimated to affect ~5,000 patients worldwide, mainly in South East Asian and Northern European populations (*11*). However, the ciliopathy shares many common and extensive abnormalities that occur during retinal development or at birth, and compromise photoreceptor structure and function. This is evident in at least four of the human ciliopathies *–CEP290, NPHP5, RPGRIP1, TULP1–* which show central outer nuclear layer (ONL) preservation with more peripheral loss, and dramatically reduced cone function which does not reflect the degree of cone cell preservation (*12*–*18*). Such extensive photoreceptor structural abnormalities, which usually involve the outer segments (OS), may preclude functional and structural correction after gene augmentation therapy, even if the transfer of a therapeutic transgene is successful. However, it is possible to examine and address these questions and facilitate the path to human treatments by using translationally relevant animal models whose disease closely parallels the human.

The canine *NPHP5* disease homologue recapitulates exactly the human disease with early onset disease, central preservation of non-functional cone photoreceptors, and relentless progression to blindness (*19*). We have used this model to examine questions of photoreceptor targeting and treatment at a stage when photoreceptors are severely diseased and active degeneration is ongoing. Treatment was delivered at a time when there is a lack of cone function because most cone OS have failed to form, and rod function was severely compromised due to marked OS structural abnormalities (*19*). Gene augmentation therapy leads to formation of normal cone OS, stabilized comparable rod structures, and immunolocalization of cell class specific markers as assessed by immunohistochemistry (IHC) and confocal microscopy. In parallel, there was robust and sustained recovery of normal retinal and visual function.

## Results

### Recovery of rod and cone function after *NPHP5* augmentation

Prior to treatment at 5-6 weeks of age, the mutant retina shows marked structural abnormalities (Fig. 1). Cones are present, but the majority lack OS based on high resolution confocal microscopy and loss of cone specific molecular markers (Fig. 1, A-C). The retina has a normal complement of rods, but the OS appear disorganized and disoriented and show extensive mislocalization of rod opsin (Fig. 1D). Despite these early structural abnormalities of the OS in both photoreceptor classes, detection of glutamylated RPGR^ORF15^ (Fig. 1E), a ciliary protein that interacts with NPHP5, suggests that the integrity of the photoreceptor ciliary transition zone is not compromised initially. At this time, TUNEL labeling shows a 10-15 fold increase in ongoing photoreceptor cell death in comparison to age-matched controls (*19*).

**Fig. 1.**
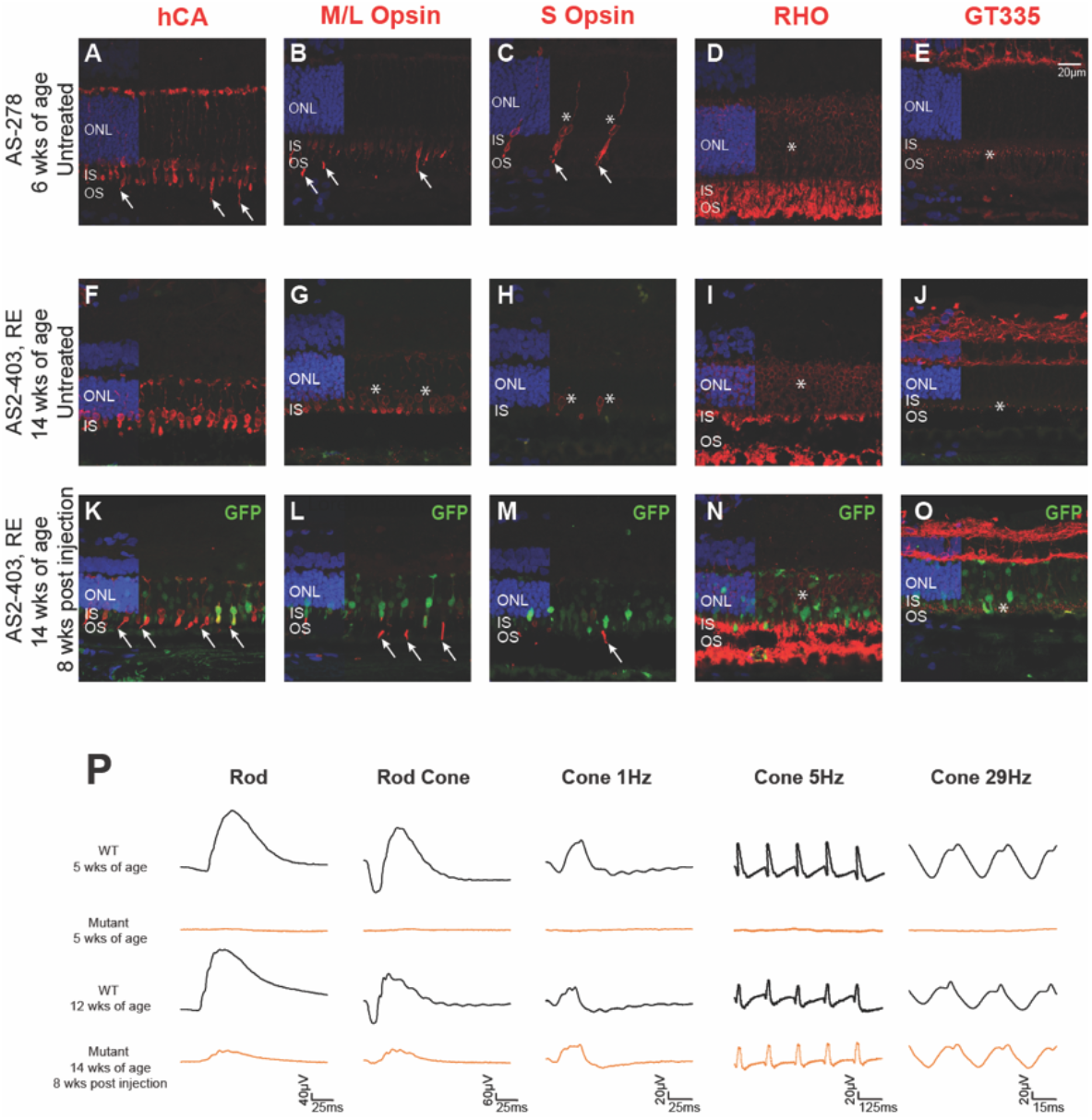
Treatment restores retinal structure and rod and cone ERG function in *NPHP5*-mutant retina. (**A to E**) Retinal structure at the 6 weeks of age time point. (**A**) Human cone arrestin (hCA) antibody shows the normal labeling of the cone IS, perinuclear cytoplasm and cone axons/pedicles, but only a small number of cone OS are present (arrows). (B, C) M/L and S cone opsins label the few OS present of the specific cone subclasses, and shows mislocalization into IS, perinuclear region and axons (*, B). (D) Rod OS are irregular and rod opsin (RHO) is mislocalized into IS and ONL (*). (E) GT335, a surrogate marker for glutamylated RPGR^ORF15^ in photoreceptor ciliary transition zone (*), shows distinct labeling at the tips of the IS. (**F-J**) Untreated regions of treated eye at 14 wks of age, 8 wks post injection (PI) shows progressive degeneration of photoreceptors, loss of ONL, and decreased labeling with GT335 in the ciliary transition zone. (**K-O**) Eye that received a subretinal injection with Vector D (scAAV2/8(Y733F)-GRK1-cNPHP5) at 1.5E+12 vg/mL; treated region identified GFP fluorescence (green). Cone-specific molecular markers show in treated areas reformation of cone OS, and reduced M/L and S opsin mislocalization. Cone OS are short and some cone IS lack an OS. The IS are more bulbous and similar in shape to untreated cone IS. Rod opsin expression in OS is intense, and mislocalization reduced (*). Localization of glutamylated RPGR^ORF15^ in the photoreceptor ciliary transition zone (*) is distinct. (**P**) Electroretinograms of the same WT control (black tracing) and mutant dogs (WM46-right eye; orange tracing) before and 8 wks after gene augmentation therapy in the mutant. There is recovery of rod and cone-mediated ERG responses with cone responses similar in waveforms and amplitudes to WT control. Light intensities used to elicit the illustrated responses are: Rod= −1.7 log cd·s·m-2; Rod Cone= 0.5 log cd·s·m-2; Cone 1Hz= 0.5 log cd·s·m-2; Cone 5Hz= 0.25log cd·s·m-2; Cone 29Hz= 0.25 log cd·s·m-2. ONL-outer nuclear layer; IS-inner segment layer; OS-outer segment layer. Hoechst nuclear label is used in all sections.

Because of the severe and rapidly progressive degenerative changes that occur early in the disease, we tested four different AAV vector constructs based on AAV2/5 and AAV2/8 with excellent photoreceptor tropism (*20*) with the aim of identifying a potential therapeutic candidate vector and promoter for treatment (Fig. 2, vectors identified as *Vectors A-D*). These vectors were AAV-based, either single stranded (ss) or self-complementary (sc); the sc vectors bypass the need for double stranded DNA synthesis prior to expression, an advantage for treating an aggressive and rapidly progressing diseases (*21*). As well, *Vector D* had a single capsid mutation, Y733F, that changed a tyrosine for phenylalanine at position 733 to bypass ubiquitination, increase nuclear targeting and transgene expression (*22*).

**Fig. 2.**
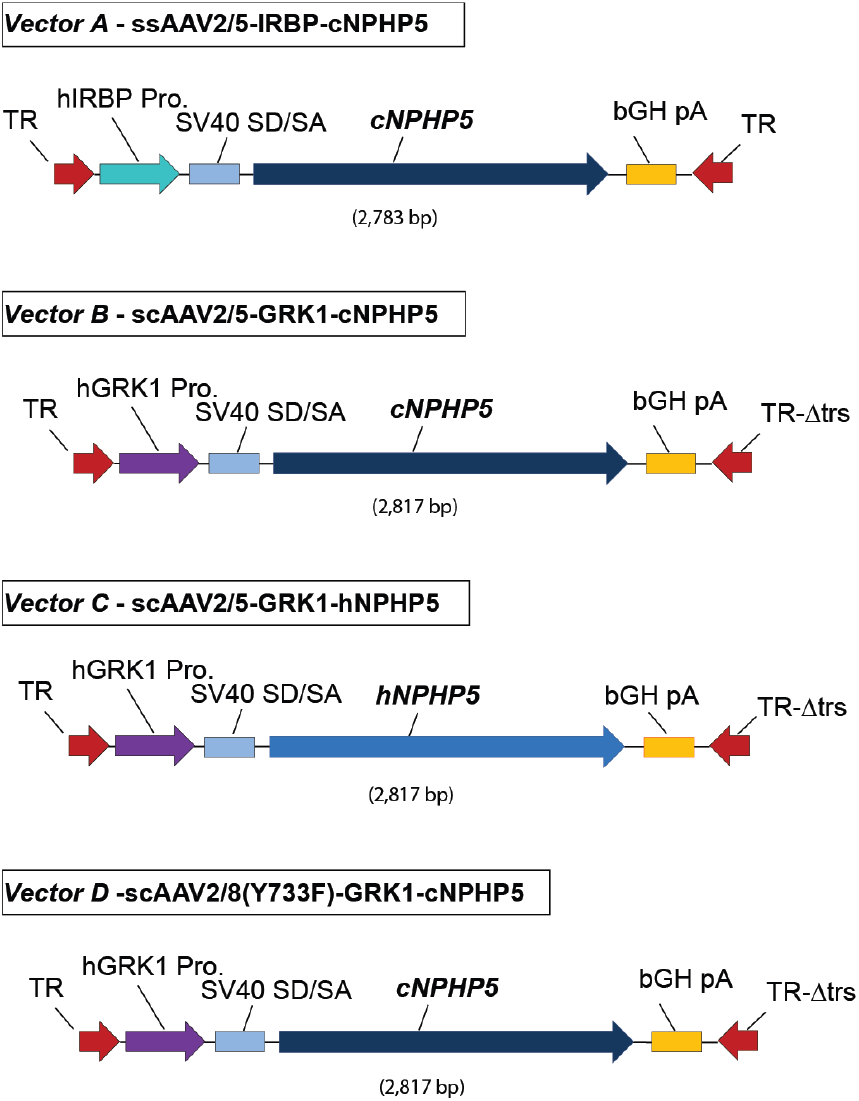
Schematics of recombinant AAV vector construct. Size (in kilo bases kb) of AAV vector cassettes containing either the human GRK1 (hGRK1) or human IRBP (hIRBP) promoters driving expression of either the canine or human *NPHP5* (*cNPHP5* or *hNPHP5*, respectively) cDNA packaged as single stranded (ss) or self-complementary (sc) constructs in either AAV8(Y377F) or AAV5 capsids. All other elements (SV40 SD/SA and poly A signals) within these constructs are identical. TR-AAV2-inverted terminal repeat; Δtrs-deletion of terminal resolution site in mutant TR; SV40 SD/SA-simian virus 40 splice donor/splice acceptor element; bGH pA-bovine growth hormone polyadenylation signal. The vectors are identified in text as *Vectors A-D*. A fifth vector, scAAV2/8(Y733F)-IRBP-*cNPHP5*, is not listed as it was used only once with no efficacy.

Early in life, *NPHP5* mutants show absence of photopic vision and cone electroretinogram (ERG) signals, and rod responses that are markedly reduced in amplitude or absent (Fig. 1P). Subretinal injection of the therapeutic vectors was directed at the high cone density temporal retinal quadrant with the visual streak/fovea-like region. The canine fovea-like region has comparable cone densities as the human and non-human primate fovea, but the presence of rods in the region makes it more comparable to the parafovea in man (*23*). Following treatment, there was a robust recovery of rod and cone ERG signals under all stimulus conditions, and the ERG cone responses were comparable to unaffected controls (Fig. 1P).

We used the four therapeutic vectors to treat 9 dogs unilaterally (n=6) or bilaterally (n=3) (table S1). When assessed 7-9 weeks post injection (PI), the 9 dogs recovered rod and cone mediated function; in these, the ERG recovery was stable for > 6 months (Fig. 3A, fig. S1). There appeared to be no apparent difference in the amplitude of the recovered responses with *Vectors B* and *C*, but the efficacy of functional recovery, although stable, was of lower magnitude in the one eye treated with a high dose of *Vector A* (Fig. 2). The single dog treated with the vector having the human transgene, *Vector C*, also had stable recovery of function, and the response amplitudes were broadly in the same range as with the other vectors. As well, there appeared to be a trend for higher amplitude responses with higher doses. The one treatment failure not included among the 9 dogs treated unilaterally or bilaterally, dog AS2-396, received scAAV2/8(Y733F)-IRBP-*cNPHP5* vector at low and high dose but did not recover ERG function at any of the 3 time points evaluated (table S1). Common to all the therapeutic vector constructs was the absence of clinical evidence of inflammation or toxicity in the 1.5E+11 to 4.7E+12 vg/mL titer range.

**Fig. 3.**
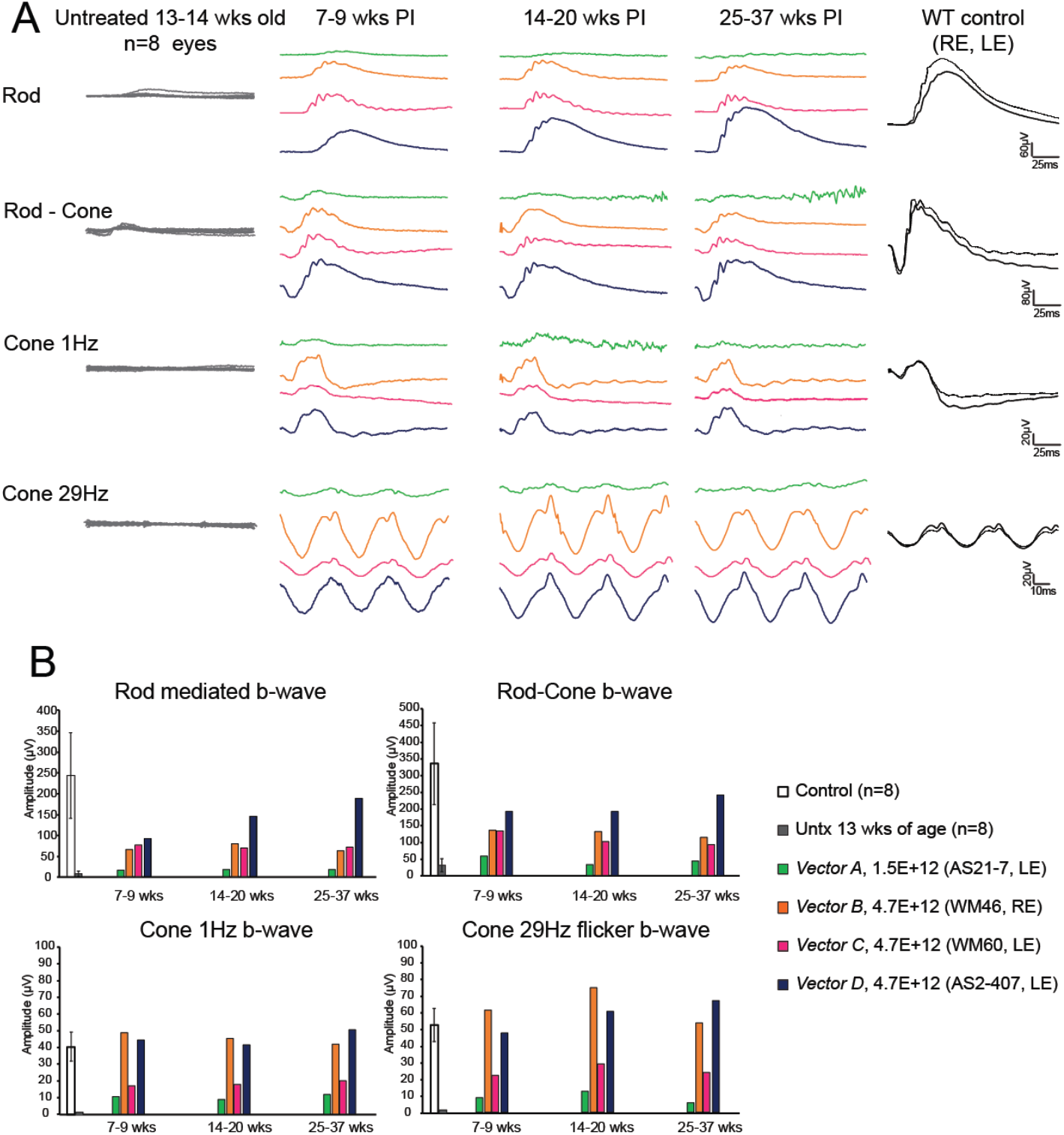
Robust and stable recovery of ERG rod and cone function following *NPHP5* gene augmentation therapy. (**A**) ERG responses from untreated mutants, 13-14 weeks of age (left column), and one wildtype (WT) control (right column) for comparison. Center panels show the individual responses of four dogs treated with 4 different vector combinations at 3 different post injection (PI) intervals. All dogs show recovery of rod and cone ERG responses that remains stable for 6 months. (**B**) Summary data of the amplitudes recorded for individual treated dogs shown in A in response to rod, rod-cone and cone selective stimuli at 3 different PI intervals. Data for control (n=4 dogs/8 eyes; 22.2+3.4 wks of age) and untreated mutants (n=8; 13-14 wks of age) is provided for comparison and represents the mean+ 1 SD of the amplitudes. Note: all but one vector has the canine *NPHP5* transgene; the human transgene is used in *Vector C* (scAAV2/5-GRK1-hNPHP5). Vector titers are in vg/mL, and eyes received 70 *μ*L subretinal injections. Individual animal identifiers are in parenthesis; LE-left eye; RE-right eye.

Suboptimal injections occurred in two of the treated left eyes (dogs AS2-404 LE and WM46 LE (table S1)), either because of a retinal tear at the retinotomy site with vector escaping into the vitreous, or failure to form a large single subretinal bleb (fig. S2*A*). In spite of the low vector volumes administered, there was distinct recovery of rod and cone mediated ERG responses that were sustained throughout the 25-32 week PI study interval (fig. S2, B and C). When compared to the optimally injected eyes (Fig. 3), the responses were lower in amplitude but distinct, and very much improved from the unrecordable ERG responses of the untreated affected dogs of comparable ages.

### *NPHP5* gene augmentation rescues vision

Functional vision was assessed in an obstacle avoidance course (movie S1) at 28 weeks PI in four unilaterally-treated mutant dogs under two ambient illumination conditions. Based on ERG functional recovery after treatment, these dogs were considered poor-moderate responders; the two high responders, dogs WM46 and AS2-407, were not included in the assessment of functional vision. All treated eyes showed faster transit time (Fig. 4A) and fewer collisions in transit (Fig. 4B), on average, under both scotopic (0.003 lux) and photopic (646 lux) conditions. The difference between treated and untreated eyes, regardless of vector, dose or transgene, were statistically significant (p<0.05), especially under photopic conditions. Under scotopic conditions, inter-animal variability was greater and significance was achieved only for 1 dog, AS21-7, that was considered a poor responder based on ERG functional recovery (Fig. 4, A and B).

**Fig. 4.**
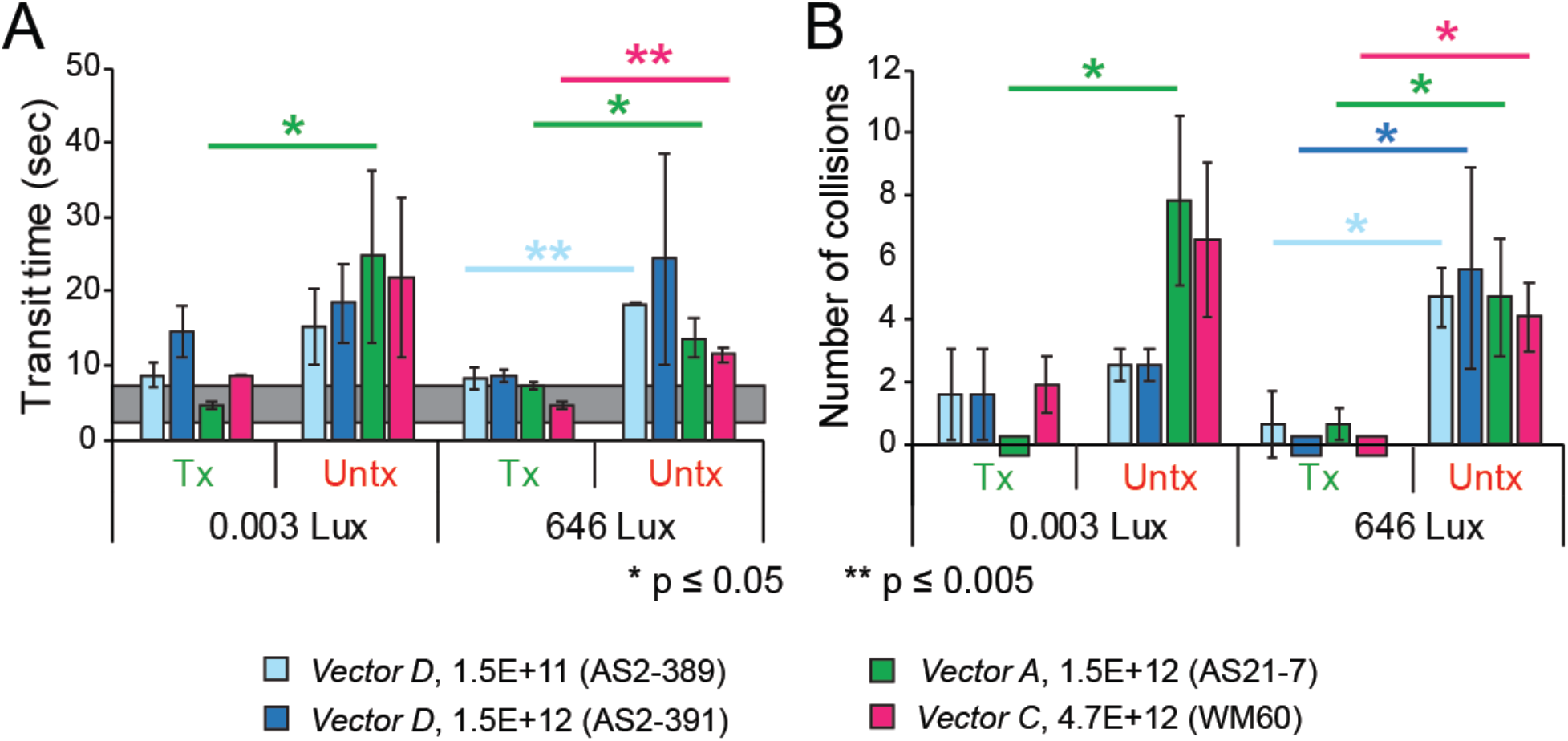
Improved visual function following gene therapy. Visual behavior results in an obstacle avoidance course for 4 *NPHP5* mutants treated unilaterally with three therapeutic vectors. Visual function was assessed under (**A**) dim scotopic (0.003 Lux) and (**B**) bright photopic (646 Lux) conditions. The horizontal gray bar in (A) represents the 95% confidence interval of the transit time for 4 WT control dogs. (B) No collisions were observed in the WT dogs. A paired t-test was used to analyze the mean difference in transit time and number of collisions under each ambient illumination between treated eyes and the contralateral control eyes, and p<0.05 was considered to be statistically significant. Tx-treated eye, Untx-untreated eye. Vector titers are in vg/mL, and eyes received 70 *μ*L subretinal injections.

### Photoreceptor layer preservation with treatment

The natural history of canine *NPHP5* disease shows widespread degeneration of the photoreceptor and ONL by ~6 months of age with a slight degree of preservation within the visual streak and fovea-like region (*19*). In comparison to wild type controls, sham injected eyes showed extensive retinal degeneration that was comparable to untreated eyes (Fig. 5, A and B). In contrast, *NPHP5* gene augmentation resulted in a distinct area of ONL preservation within the treated region the corresponded with the recovery of retinal and visual function (Fig. 5, C-F and fig. S3, A-D). Comparable results were obtained with *Vectors A-D*, and there were no obvious differences that could be attributed to dose or the canine vs human *NPHP5* transgene. The distribution of ONL rescue, however, was dependent generally on the bleb size (Fig. 5, C-F and fig. S3, A-D). In the one dog where the focal bleb became widely distributed throughout the superior retinal quadrants because of an entrapped air bubble in the injection fluid, ONL preservation was commensurate with the vector distribution (Fig. 5F); the untreated fellow eye showed advanced retinal degeneration.

**Fig. 5.**
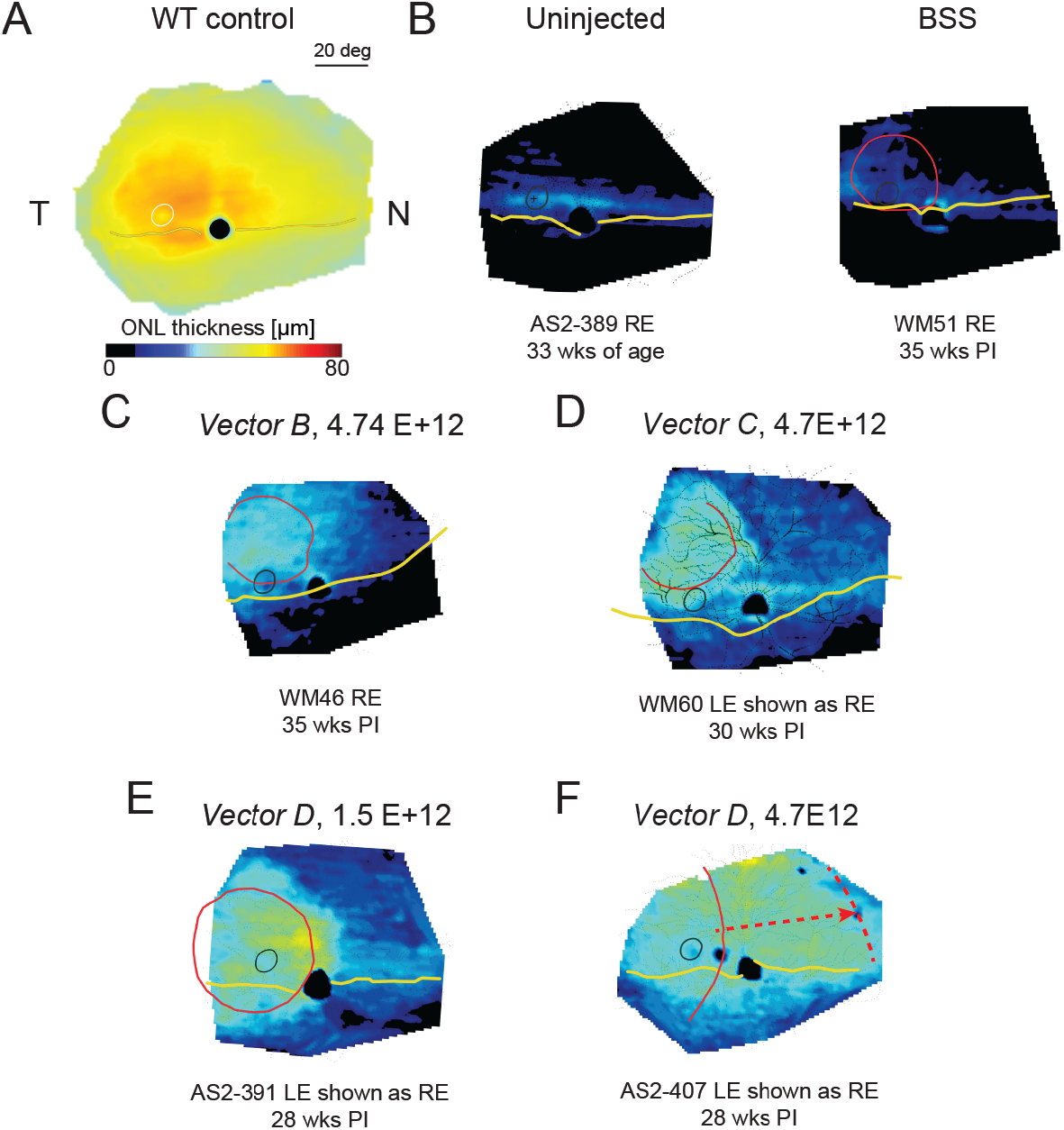
In vivo retinal imaging shows preservation of photoreceptor outer nuclear layer (ONL) in treated areas. Topographical maps of ONL thickness in (**A**) wildtype (WT) control and (**B**) untreated or sham treated eyes. All eyes treated with the therapeutic vectors (**C-F**) show preservation of ONL thickness in comparison to untreated areas of the same eye. Retained ONL thickness in some cases extends beyond the bleb border into a penumbral region. Anatomic landmarks added include the fovea-like region (temporal white or black circle), the optic disc (black), and the tapetal boundary (wavy yellow line). In treated eyes, the red line identifies the subretinal injection bleb determined photographically immediately after the injection. The dashed arrow/red line in F shows the putative spread of the subretinal injection bleb from a trapped air bubble in the injection fluid. BSS-balanced salt solution; LE-left eye; RE-right eye; PI-post injection interval. Vector titers are in vg/mL, and eyes received 70 *μ*L subretinal injections.

### Expression of therapeutic transgene in treated retinas

Neither commercial nor custom made antibodies directed against the N- or C-terminus of human NPHP5 gave specific immunolabeling with or without antigen retrieval when used on retinas treated with the canine NPHP5 transgene (table S2). To specifically localize *NPHP5* transgene expression we used RNA *in situ* hybridization (ISH). Following gene therapy, the treated areas showed intense RNA hybridization signals primarily localized in the ONL, with scattered signals present in the inner retinal layers and RPE, albeit at much reduced in intensity (fig. S4, A to C). In contrast, the untreated areas of the same eye, or the uninjected fellow eye, showed only very slight background signal. Co-injection with an AAV2/5-IRBP-*GFP* reporter vector, and detection of GFP fluorescence in similar retinal areas on sequential sections confirmed that *NPHP5* mRNA expression was limited to the subretinally-treated region. Similar ISH labeling patterns were found in dogs treated with *Vector D* and analyzed at 8, 27 and 29 wks PI. At the longer PI intervals, the treated areas had much better ONL preservation than untreated areas of the same or fellow eye, and GFP fluorescence showed that the treated photoreceptor IS were now elongated and had a normal silhouette (fig. S4, B and C, lower panels).

### Reforming normal photoreceptor structure after gene augmentation

Immunohistochemistry was used to examine the photoreceptor structural correlates that accompanied the recovery of retinal function following gene therapy. Eight weeks after subretinal injection, the treatment areas delineated by GFP fluorescence show that cones began to reform. Cone OS develop, start to elongate and express M/L- and S-cone opsins. The cone IS were short and bulbous, similar to those in untreated areas, but cone opsin mislocalization was reduced (Compare Fig.1, F-H and K-M). Similarly, rod opsin mislocalization was minimized, and RHO immunolabeling was more intense (Fig. 1, I and N). At the photoreceptor ciliary transition zone, there was more intense expression of glutamylated RPGR^ORF15^, a protein that interacts with NPHP5 in photoreceptors (*4*, *24*), than in untreated areas (Fig. 1, J and O).

Longer PI intervals clearly showed the significant positive treatment effects. Cones IS became slender and longer, and, with few exceptions, all expressed M/L- or S-cone opsins in well formed elongated OS (Fig. 6, A-C and F-H; fig. S5, A-C and F-H). Similarly, based on high resolution confocal microscopy and IHC, rod IS and OS structure were normal, and glutamylated RPGR^ORF15^ localization in the photoreceptor ciliary transition zone was also normal (Fig. 6, D and E and I and J; fig. S5, D and E and I and J). In contrast, untreated areas of the same eye showed marked thinning of the ONL, loss of photoreceptors, and extensive mislocalization of rod and cone opsins into the perinuclear region of the few remaining ONL nuclei. These positive treatment effects emphasized that very abnormally developed photoreceptors can reform normal structure following gene augmentation therapy, and that this reversal was stable.

**Fig. 6.**
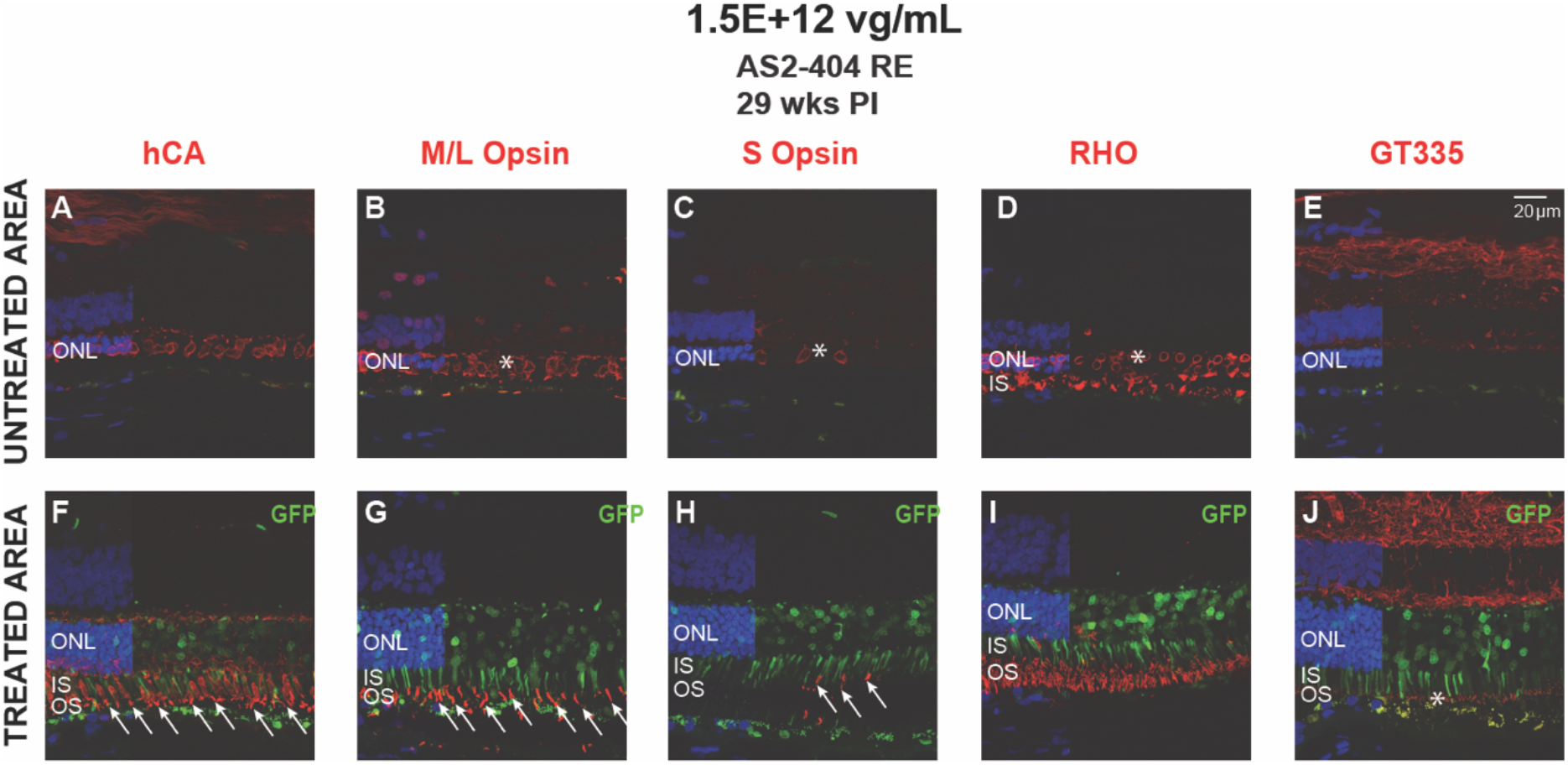
Stable recovery of normal rod and cone structure after treatment. (**A** to **E**) Sections from untreated and (**F** to **J**) treated areas (*Vector D*; scAAV2/8(Y733F) GRK1-*cNPHP5*) of the right eye of dog AS2-404 at 29 weeks following therapy; treated areas identified by GFP fluorescence (green) in photoreceptor and outer nuclear layer nuclei. (A to C and F to H) Cone specific molecular markers show normal cone structure in treated areas, and all cone IS have corresponding OS (white arrows). Untreated areas show absence of cone IS and OS, and cone opsin mislocalization to ONL (B and C*). (D and I) Rod OS are structurally normal, and opsin mislocalization is almost completely reversed in treated area, but in untreated area rod degeneration has advanced and remaining rod nuclei have perinuclear RHO localization (D*). (E and J) Localization of glutamylated *RPGR^ORF15^* in the ciliary transition zone (*) is distinct, while the untreated region has none. Hoechst nuclear label is used in all sections. ONL-outer nuclear layer; IS-inner segment; OS-outer segment; RE-right eye; vg/mL-vector genomes/mL.

## Discussion

The momentum generated by FDA’s approval of LUXTURNA™ (voretigene neparvovec-rzyl) for treating patients with biallelic *RPE65* mutations (*25*, *26*) has given hope that successful treatment for previously untreatable blinding diseases is possible. Gene therapy trials for other inherited retinal diseases have been initiated (*27*), and provide further optimism for eventual cures. Among candidate diseases, the ciliopathy disorders provide a compelling therapeutic target, especially those for which there is a dissociation of structure and function. The absence of retinal function measured objectively by ERG, decreased retinal sensitivity and/or impaired vision performance in a mobility test permits an early assessment of a successful functional outcome in treated eyes in which non-invasive analysis ascertains the presence of a sufficient number of photoreceptors suitable for treatment (*28*, *29*). These outcome measures can be identified soon after treatment, as was the case in the *RPE65* clinical trials (see (*30*, *31*) for review and study summaries), a form of LCA also characterized by a dissociation of structure and function (*28*). In contrast, outcomes for therapies that aim to halt disease progression and preserve vision are more difficult to assess quickly, and usually require many years to establish positive outcomes (*32*).

The canine *NPHP5* model is an excellent platform to develop therapies for the class of ciliopathies characterized by early onset, aggressive clinical course, and dissociation of retinal structure and function. In both dogs and patients there is marked peripheral rod loss, but the degree of cone function loss does not reflect the degree of central cone preservation. In patients, cross-sectional *in vivo* imaging of the cone-only foveal region shows that in most the ONL thickness is normal, and the presence of a distinct ellipsoid region in the foveal cones (EZ line) indicates that cone IS have formed; similarly, dogs show that cones, both in the cone enriched fovea-like region and more peripherally, initially have normal numbers, but these lack an outer segment (*19*).

In dogs, the disease is characterized by an overlap of abnormal photoreceptor development and progressive degeneration occurring in two phases of photoreceptor loss, an early peak at ~5 wks of age and a constant rate of loss thereafter (*19*). At the initial disease stage, ~85% of cones, but not rods, fail to form an OS; however, the rod OS have irregular contours and there is extensive rod opsin mislocalization into the IS, ONL and synaptic terminals. In spite of these cone structural abnormalities, a subset of cone-specific genes, particularly the cone opsins and the cyclic nucleotide gated a and β subunits, are unchanged in expression while the expression of rod-specific and rod/cone-specific genes are downregulated (*19*). Thus the structural, functional and molecular abnormalities in this form of LCA result in severely compromised photoreceptors in early life.

Regardless of the four AAV constructs used, gene augmentation therapy delivered at 5-6 weeks of age resulted in robust recovery of rod- and cone-mediated ERG responses when evaluated 7-9 wks PI, and remained stable to the end of the study. The responses showed normal waveforms, and recovery of cone responses was dramatic as there was complete absence of cone signals at the time of treatment because most of the OS had failed to form (present study and (*19*)). For functional recovery to occur, the cones had to reform the OS and express and localize the phototransduction components in the correct subcellular compartment. Given that the subset of cone-specific genes analyzed using a canine-specific qRT-PCR profiling array showed normal expression, it is likely that expression of wild type (WT) NPHP5 in cones restored normal trafficking of proteins through the cone ciliary transition zone resulting in OS formation and restoration of function (*19*).

Rods too have severe functional deficits that result from compromised structure. At the time of treatment rod OS are present but their contours are irregular and their parallel orientation is lost, there is extensive rod opsin mislocalization into the IS and ONL, and rod cell death is at its peak (*19*). Rod-isolated ERG responses could be present, but reduced in amplitude by 60-75%, or absent (present study and (*19*)). Impaired trafficking of key phototransduction proteins through the ciliary transition zone explains both opsin mislocalization and abnormal rod ERG function prior to therapy. Following gene augmentation therapy, rod function is restored and stabilized during the PI period studied.

In this study, we tested the safety, efficacy and dose ranges of four therapeutic vector combinations with the canine or human *NPHP5* transgenes regulated by either IRBP or GRK1 promoters and packaged either as single stranded or self-complementary DNA constructs (Fig. 2). Our initial aim was to identify a superior vector for translational applications. However, we found that a positive therapeutic response is not limited to a specific vector, promoter or transgene combination, but that the mutant retina appeared quite promiscuous in responding positively to therapeutic intervention. In general, the principal determinants of a positive response appeared to be the dose, and the size/distribution of the subretinally injected vector.

At the doses used, the four vectors were well tolerated with a good safety profile, and resulted in stable recovery of cone and rod function. We found no apparent differences in the functional recovery with either the scAAV2/5 and scAAV2/8(Y733F) vector pseudotypes containing the GRK1 promoter. However, the small sample size and multiple vector/promoter/transgene combinations used precluded definitive assessment of relative efficacy. There appeared to be a trend for higher amplitude responses with increasing doses. Again, the small sample size precluded statistical analysis. Lastly, a fifth vector, scAAV2/8(Y733F) IRBP-cNPHP5, was tested in two eyes of a single animal, but failed to restore function and only partially reformed photoreceptor structure, yet stopped ONL loss in the one treated retina.

There were differences, however, in the degree of recovery of rod and cone function following treatment. Considering that the 70 *μ*L injection volume covered no more than ~25% of the retinal surface, the amplitude of the recovered rod b-wave responses were commensurate with the treatment area, approximately 20-30% of normal control values depending on the stimulus conditions used. The one animal where the injection bleb extended into the rod enriched nasal superior quadrant, and thus covered ~40-50% of the retinal surface, showed rod ERG responses close to control values.

Cone responses, on the other hand, proportionally were higher in amplitude, and at the 1.5 and 4.7E+12 vg/mL titers with the canine transgene, were comparable to WT controls; the amplitudes of the recovered cone responses remained unchanged during the 6 month PI period (Fig. 3B). As the treatment was directed to the cone enriched temporal quadrant that included the visual streak and the fovea-like region, it is likely that a higher cone population density was targeted than if treatments had been directed to the nasal or inferior retinal quadrants where cone densities are lower (*23*, *33*).

Like in humans, functional vision is severely compromised in *NPHP5* mutant dogs early in life, particularly under photopic conditions. This clinical observation was supported by objective vision testing in an obstacle avoidance course. Following treatment, transit times were decreased as did the number of collisions. As it takes several weeks of training after dogs have achieved physical maturity, the improvement of visual function could not be evaluated at earlier time points after treatment, but only at the end of the current study. At that time, there was a definite improvement in scotopic and photopic vision that was neither AAV capsid, promoter, dose, nor transgene-specific as the three vector combinations tested gave comparable results.

Underpinning the recovery of retinal function and vision were events occurring at the level of the photoreceptor cells that ensured the stability of the therapeutic intervention. Based on in situ hybridization results, treated regions showed robust expression of the therapeutic transgene; expression was present at the earliest time point analyzed, 8 weeks PI, and was qualitatively unchanged at the 27-29 wk PI time points. As the vector was administered by subretinal injection, it is not surprising that some cells that border the subretinal space – RPE, photoreceptors and Müller cells – were also transduced. However, it was not expected that they would express the message, even at low levels, as the promoters used were photoreceptor-specific. Given that these promoters are shortened versions of the natural promoters, it is likely that targeting specificity was limited, and leaky expression in other cell classes occurred. Whether the transduced cells express the protein is presently unknown as specific antibodies that can detect canine NPHP5 are not available. What is clear, however, is that there is no apparent damage to any of the cells that express the *NPHP5* transcript.

Within eight weeks of treatment, and in parallel with recovery of rod and cone ERG function, photoreceptors in the treated area began to reform normal structure. Initially, shortened OS were associated with many of the cone IS which still had the bulbous contours characteristic of the untreated mutant retinas. The newly formed OS expressed M/L- and S-cone opsins, and reversed the protein mislocalization (Fig. 1 and (*19*)). Rods too showed more robust rod opsin expression that accompanied a partial reversal of this mislocalized protein. Our prior studies indicated the presence of normal ciliary transition zone based on MAP9 expression in mutant retinas; now we show that treatment maintains expression of glutamylated RPGR^ORF15^, an NPHP5 interacting protein also localized in the ciliary transition zone (*4*, *24*). The reversal of structural abnormalities in the treated retinas was stable. By the end of the study, ~27-29 weeks PI, the rods and cones in treated areas had regained normal structure, and expressed key proteins characteristic of each photoreceptor class. It is important to note, however, that normal structural assessment was based on high resolution confocal microscopy and IHC, and not electron microscopy which would have been required to critically assess the internal structure of the photoreceptor OS.

Along with the stable recovery of photoreceptor structure was the preservation of the ONL. At the ~5-6 weeks of age when treatment was initiated, photoreceptor apoptosis is at its peak and subsequently declines to more constant rates (*19*). It is likely that many of the cells that were successfully transduced by treatment were already committed to the cell death pathway and would have died regardless of therapy (*34*). This would have resulted in a slight thinning of the ONL in the post treatment period in spite of a positive treatment effect. As well, the canine retina, especially the ONL, thins with aging in the postnatal period. This process begins 4 weeks after birth, and decays exponentially until ~25 wks of age as the globe increases in axial length (*35*, *36*). Assessment of successful preservation of photoreceptor ONL needs to take into account this age-associated thinning and determine successful outcomes not on the ONL thickness at the time of treatment, but on the expected thickness after 25 wks of age (see Fig. 5 in (*36*)). By these measures, it is clear that treatment arrested disease progression and ONL thickness stabilized.

From the *en face* topographic maps of ONL thickness, it was clear that the region of treatment efficacy extended beyond the bleb borders that were photographically documented at the time of subretinal injection, and formed what is termed a penumbral region (*37*). The *en face* maps showed a variability in the extent of the penumbral region. In some, preservation was restricted to a narrow region surrounding the treatment bleb, while in others, the preservation was more widespread (compare Fig, 5E (AS2-391) with fig. S3*B* (WM51 LE)). The rescue effect outside of the treatment area has been observed in other studies, e.g. (*37*, *38*), and is partly attributed to expansion of the subretinal bleb prior to reattachment (*39*).

Like the treated *NPHP5* mutant dogs, gene augmentation of knock out (KO) mice that have an artificial all S-cone environment by being double homozygous for *Nphp5^-/-^;Nr^-/-^* also shows rescued photopic function, the treated cones form axonemes, and the phototransduction proteins relocate to the OS (*40*). While it is not clear that reforming cone OS in an artificial all S-cone environment is relevant for human applications, it provides further support that cones have the ability to reform OS that never developed, or were lost through disease (*41*), and emphasizes the plasticity of the photoreceptor sensory cilia that continually renew their OS disc membranes (*42*). Here we showed that cone OS recovery occurs in a normal rod and cone enviroment that models closely the human retina, and its disease phenotype. From the perspective of translational applications, such a model that is a disease homolog, and recapitulates the molecular and cellular defects along with phenotypic identity, will facilitate progress in the clinical development continuum. Furthermore, it will inform on therapeutic efforts for the other ciliopathies that show similar central ONL preservation with more peripheral loss, and dramatically reduced cone function which does not reflect the degree of preservation of cone nuclei (*12*–*17*).

## Materials and Methods

### Study design

The objective of this study was to determine if this aggressive and severe ciliopathy would show a positive response to AAV mediated gene augmentation therapy, and to identify the potential therapeutic candidate vector and promoter for translational development. To this end, mutant dogs were produced by breeding mutation identified affected dogs maintained in a closed research colony. The mutant dogs were subretinally injected with one of four AAV vector constructs carrying a full-length canine or human *NPHP5* cDNA under the control of either human IRBP or GRK1 promoters. Assessment of the response to gene transfer and stability of treatment was made by means of longitudinal analyses using clinical ophthalmic examinations, *en face* and cross sectional in vivo retinal imaging, electroretinography, psychophysical vision testing, and, at the end of study, morphological and *in situ* hybridization evaluation of retinal sections. Methodological details are provided in Supplementary Material, Methods.

## Supporting information

NPHP5 Supp Material

## List of Supplementary Materials

Materials and Methods

Fig. S1. Robust and stable recovery of ERG rod and cone function following *NPHP5* gene augmentation therapy.

Fig. S2. Stable functional recovery even with suboptimal treatments.

Fig. S3. In vivo retinal imaging shows preservation of photoreceptor outer nuclear layer (ONL) in treated areas.

Fig. S4. Therapeutic *NPHP5* transgene is expressed in treated area of injected eyes.

Fig. S5. Stable recovery of normal rod and cone structure after treatment.

Movie S1: Restoration of Visual Function in Canine NPHP5 LCA by Gene Augmentation Therapy.

Tables S1 and S2

## Acknowledgements

The authors thank Dr. John E. Dowling of Harvard University for critical review of the manuscript and helpful comments, Vince Chiodo of the University of Florida for vector production, Svetlana Savina, Natalia Dolgova and Evelyn Santana for technical support, Sommer Ainsworth for vision behavior testing and Dr. Karolina Roszak for carrying out some of the *in vivo* studies-OCT, ERG, and preparations for the subretinal injections. Terry Jordan and the staff of RDSF for excellent animal care and support of in vivo studies, and Lydia Melnyk for research coordination.

## Author contribution

GDA designed and implemented the studies, analyzed the data, wrote original and final draft, and supervised the research activity. AVC analyzed and interpreted the noninvasive imaging results, provided critical review and advice at different stages of the study, and was involved in the review and editing of the manuscript. VLD carried out different aspects on the in vivo studies, reviewed and wrote parts of the manuscript, and produced most of the figures. ARG and SI carried out eye examinations, medical assessments, ERG and OCT studies. RS carried out the RNA *in situ* hybridization studies with the assistance of RN. MS used OCT imaging data to create the en face maps of ONL thickness. SLB and WWH designed and produced the vectors. SGJ provided critical review and advice at different stages of the study, and was involved in the review and editing of the original draft of the manuscript. WAB designed and implemented the studies, carried out the subretinal injections, assessed retinal structural outcomes by histology and immunohistochemistry, analyzed the data, wrote parts of the original and final draft, and supervised the research activity

