## Supplementary material for "Gene Therapy Reforms Photoreceptor Structure and Restores Vision in *NPHP5*-associated Leber Congenital Amaurosis": NPHP5 Supp Material

**This document includes:**

Materials and Methods
Fig. S1. Robust and stable recovery of ERG rod and cone function following *NPHP5* gene augmentation therapy.
Fig. S2. Stable functional recovery even with suboptimal treatments.
Fig. S3. In vivo retinal imaging shows preservation of photoreceptor outer nuclear layer (ONL) in treated areas.
Fig. S4. Therapeutic *NPHP5* transgene is expressed in treated area of injected eyes.
Fig. S5. Stable recovery of normal rod and cone structure after treatment.
Movie S1: Restoration of Visual Function in Canine NPHP5 LCA by Gene Augmentation Therapy.
Tables S1 and S2

### Supplementary Methods

CANINE *NPHP5* MODEL

The *NPHP5* model originates from a research colony used for mutation identification (*43*) and disease characterization; affected dogs have a severe early-onset retinal disease, but no associated renal abnormalities (*19*). They were produced by mating parents of known genotype to produce affected progeny. Ten (males=2; females=8; 14 eyes) dogs were used for the gene therapy studies (table S1). As non-affected controls for ERG studies, both eyes of 5 dogs (males=4, females=1; 10 eyes) were used; these were heterozygous for mutations in other autosomal recessive retinal disease genes (*PDE6B* (n=1 (*44*)), *CNGA3* (n=1 (*45*)), *STK38L* (n=3 (*46*)) where the carrier animals are indistinguishable from normal in terms of retinal structure and function. ERGs were done in these control dogs at 22.2±3.4 wks of age. Another WT male control dog had an ERG performed at 5 and 12 wks of age. All dogs were bred and maintained at the Retinal Disease Studies Facility, Kennett Square, PA, and housed under identical conditions of diet, medications, vaccinations, and ambient illumination with cyclic 12 hrs ON-12 hrs OFF. For terminal procedures, the dogs were euthanatized with an overdose of Euthasol euthanasia solution (Virbac), and the eyes immediately enucleated. The research was conducted in full compliance with the University of Pennsylvania Institutional Animal Care and Use Committee (IACUC) approval, adhered to the Association for Research in Vision and Ophthalmology (ARVO) Resolution for the Use of Animals in Ophthalmic and Vision Research, and followed the recommendations in the Guide for the Care and Use of Laboratory Animals of the National Institutes of Health.

VIRAL VECTORS

Four different therapeutic adeno-associated viral (AAV) vector constructs were developed; these are identified in the main text as *Vectors A-D* (Fig. 2). They carried either the canine (c) or human (h) full length *NPHP5* cDNAs that were synthesized based on the wildtype canine (*43*) or human (Genbank, accession number AY714228.1) sequences, and incorporated a canonical Kozak sequence at the translational initiation site (Genscript). *NPHP5* transgenes were placed under control of one of two photoreceptor specific promoters: a 292 nt segment (positions 1793–2087) of the human G-protein-coupled receptor protein kinase 1 promoter (hGRK1, GenBank AY327580), or a 235 nt segment of the human interphotoreceptor retinoid-binding protein (IRBP) promoter that includes the cis-acting element identified in the mouse proximal promoter (*47-49*). Single stranded (ss) or self-complementary (sc) cassettes were packaged into one of two capsids: AAV8(Y733F) or AAV5. A fifth vector, scAAV2/8(Y733F)-IRBP-*cNPHP5*, is not included in Fig. 2 and was only tested in both eyes of one dog and did not result in recovery of retinal function or preservation of photoreceptor structure.

AAV vector preparations were produced by triple plasmid co-transfection using PEI (*50*). Vector purification involved discontinuous iodixanol step gradient followed by ion-exchange column chromatography (HiTrap Q Sepharose column) as previously described (*51*). The vector-containing fractions eluted from the columns were concentrated, and titered for DNase-resistant vector genomes by RT-PCR relative to a standard. Finally, the purity of the vector was validated by silver-stained SDS–PAGE and vector preparation confirmed to have endotoxin levels <5.0 EU/mL.

VECTOR ADMINISTRATION AND POST INJECTION TREATMENT

Subretinal injections of recombinant AAV vectors were performed under general anesthesia in mutant dogs between 5.1 and 5.7 wks of age as previously described (*49, 52, 53*) (Supplemental Table 1 details the animals, vectors and promoters used). 70 $\mu$L of viral vector diluted in balanced salt solution (BSS) or vehicle (BSS) were delivered using a transvitreal approach without vitrectomy with a custom-modified RetinaJect subretinal injector (SurModics) (*54*) under direct visualization with an operating microscope; anterior chamber paracentesis was performed immediately post injection (PI) to prevent increase in intraocular pressure. Directly after injection, location of the subretinal bleb was documented by fundus photography (RetCam Shuttle). The injections were directed to the temporal quadrant and were considered optimal when a large uniform bleb formed that included the area centralis and fovea-like region of the retina even when the injection was slightly off-center (*23*). Two eyes had suboptimal injections where the bleb size was small, flat or multiloculated, or there was a visible retinal tear at the retinotomy site (fig. S2). In all cases, the injection bleb flattened and the retina reattached within 24 to 48 hr PI. Ophthalmic examinations, including biomicroscopy, indirect ophthalmoscopy, and fundus photography, were conducted at 24 hrs, 48 hrs, and 5 days PI, and then weekly for the first 2 months followed by a monthly eye examination thereafter throughout the injection–end point evaluation time interval.

Postoperative treatment was identical to that used in previous studies (*49, 53*) and described in detail below: on the morning of surgery, dogs received oral (amoxicillin trihydrate/clavulanate potassium 12.5-20 mg/kg) and topical antibiotic (gentamicin sulfate 0.3% solution; 1 drop/eye) and anti-inflammatory (oral prednisone 1mg/kg; topical prednisolone acetate 1% suspension, 1 drop/injected eye; topical flurbiprofen 0.03%, 1drop/injected eye) medication and topical mydriatics (tropicamide 1% and phenylephrine 10%, 1 drop/injected eye 3 times, 30 minutes apart). Immediately after the end of the surgical procedure, a subconjunctival injection of 4 mg of triamcinolone acetonide (Kenalog 40 mg/mL), and topical antibiotic + steroid ointment (NeoPolyDex ointment ¼ inch strip/eye) was applied to each injected eye. In the afternoon of the procedure, the animals received a second dose of oral antibiotics (amoxicillin trihydrate/clavulanate potassium 12.5-20 mg/kg) and oral anti-inflammatory (prednisone 1 mg/kg) medication. Topical application of atropine sulfate 1% was given once daily (1 drop/injected eye) for 1 week. Topical application of corticosteroid suspension (prednisolone acetate 1% suspension 1 drop/injected eye) was applied twice a day in the injected eyes for 2 weeks and then once daily for the following 2 weeks. Oral administration of antibiotics (Amoxicillin trihydrate/clavulanate potassium 12.5-20 mg/kg) was given twice a day for 5 weeks. Oral administration of corticosteroids (Prednisone) was given twice a day for 2 weeks at the dose of 1 mg/kg, then once a day for 2 weeks at the dose of 1 mg/kg, and was then discontinued. At 4 weeks post-injection a second subconjunctival injection of 4 mg of triamcinolone acetonide (Kenalog 40 mg/mL), was given in the injected eyes after topical anesthesia (Proparacaine 0.5%, 1 drop/eye). Dogs were not tested for neutralizing antibodies directed against the capsid protein prior to subretinal injection of the AAVs.

### Electroretinography

Full-field flash ERGs were recorded with an Espion ERG system (Diagnosys) under general anesthesia (induction with IV propofol; maintenance with isofluorane) using a custom-built Ganzfeld dome fitted with the LED stimuli of a ColorDome stimulator (Diagnosys) and methods previously described (*53*). Waveforms were processed with a digital low-pass 50 Hz filter to reduce recording noise if necessary. After 20 min of dark adaptation, rod and mixed rod-cone–mediated responses (averaged four times) to single 4-ms white flash stimuli of increasing intensities (from −3.74 to 0.5 log cd·s·m−2) were recorded. Following 5 min of white light adaptation (1.025 log cd·m−2), cone-mediated signals (averaged 10 times) to a series of single flashes (from −2.74 to 0.5 log cd·s·m−2) and to 5-Hz (averaged 20 times; from −2.74 to 0.25 log cd·s·m−2) and 29.4-Hz flicker (averaged 20 times; from −2.74 to 0.25 log cd·s·m−2) stimuli were recorded. These protocols separately assessed rod- and cone-mediated responses.

#### Optical Coherence Tomography

*En face* and retinal cross-sectional imaging was performed with the dogs under general anesthesia (induction with IV propofol; maintenance with isofluorane) as previously reported (*53*). Overlapping *en face* images of reflectivity with near-infrared illumination (820 nm) were obtained (Spectralis HRA+OCT) with 30° and 55° diameter lenses to delineate fundus features such as optic nerve, retinal blood vessels, boundaries of injection blebs. Custom MatLab 7.5 programs (MathWorks) were used to digitally stitch individual photos into a retina-wide panorama. Spectral domain-optical coherence tomography (SD-OCT) was performed with linear and raster scans (Spectralis HRA+OCT). Overlapping 30° X 20° raster scans were recorded covering large regions of the retina. Post-acquisition processing of OCT data was performed with custom programs (MatLab 7.5). For retina-wide topographic analysis, integrated backscatter intensity of each raster scan was used to locate its precise location and orientation relative to retinal features visible on the retina-wide mosaic formed by NIR reflectance images. Individual longitudinal reflectivity profiles (LRPs) forming all registered raster scans were allotted to regularly spaced bins (1° X 1°) in a rectangular coordinate system centered at the optic nerve;

LRPs in each bin were aligned and averaged. Intraretinal peaks and boundaries corresponding to the ONL were segmented using both intensity and slope information of backscatter signal along each LRP. For all topographic results, locations of blood vessels, optic nerve head, and bleb boundaries were overlaid for reference.

#### VISUAL BEHAVIOR-OBSTACLE AVOIDANCE COURSE

The methods to evaluate vision in dogs in an obstacle avoidance course have been previously reported (*53, 55*). For this study we used an abbreviated protocol that evaluated visual function under two ambient light conditions: very dim scotopic (0.003 lux) and bright photopic (646 lux) illumination. Eyes were tested individually by placing an Aestek opaque corneo-scleral shield (Oculo-plastik Inc.) over each cornea after topical anesthesia (proparacaine 0.5%; 1 drop/eye). Both eyes were tested three times under each light intensity, and the position of the 5 panels was randomly changed between each of the 3 trials/eye/illumination. The contralateral eye was tested with the same set of panel positions. Random selection of the eye to be tested was made before the session. Dogs were first adapted for 20 min to the lowest ambient illumination (0.003 lux) before running through the course; subsequently the room illumination was increased to the photopic level of brightness (646 lux) and dogs adapted for 10 min and tested as previously described. Two digital Sony Handycam cameras (Sony) located above the obstacle course recorded the navigation performance of the dogs. The infrared imaging function of the camera enabled recording under the dimmest light conditions. An experienced observer who was masked to the experimental design reviewed all the videos to measure for each trial the transit time in seconds between the first forward motion at the entrance of the course and the moment the animal completely passed through the exit gate, and the total number of collisions. A paired t-test was used to analyze the mean difference in transit time and number of collisions under each ambient illumination between treated eyes and the contralateral control eyes, and $p<0.05$ was considered to be statistically significant.

#### RETINAL MORPHOLOGY, IMMUNOHISTOCHEMISTRY (IHC) AND IN SITU HYBRIDIZATION

Eyes were immediately enucleated after euthanasia, fixed in 4% paraformaldehyde (PFA) for 3 hrs followed by 2% PFA for 24 hrs, trimmed, cryoprotected, and embedded in optimal cutting temperature media as previously reported (*49, 53*). Preparation of all solutions was carried out with DEPC-treated water, and all tissue handling used sterilized instruments and gloves to minimize RNase contamination. Retinal cryosections (10-12 $\mu$m thickness) were rinsed in phosphate-buffered saline (PBS) with 0.5% Triton X-100 (PBST), and incubated in blocking solution [4.5% Cold Water Fish Gelatin (G7765, Sigma-Aldrich)/5% Bovine Serum Albumin (A7906, Sigma-Aldrich)/0.05% Sodium azide (S8032 Sigma-Aldrich) in PBST] for 20 min. Retinal sections were then incubated with primary antibodies diluted in blocking solution at 4°C overnight in a slide rack for immunostaining (10098889; Thermo Scientific). Several commercial and custom-made antibodies against human NPHP5 were used, but gave no specific immunolabeling ((*56*) and table S2). The sections were rinsed in PBS for 5 min 3 times, and then incubated with the secondary antibodies (Alexa Fluor Dyes, Thermo Fisher Scientific) diluted (1:200) in blocking solution at room temperature for one hour. Sections were then rinsed for 5 min 3 times. Finally, sections were incubated in Hoechst nuclear stain (Hoechst 33342, Invitrogen) diluted (1:100) in PBS for 30 min at room temperature. Slides were mounted with coverslips with Fluoromount G (0100-01, Southern Biotech). TUNEL labeling was done with the *in situ* cell detection kit (Roche). H&E stained sections were examined by brightfield microscopy and single or double immunolabeled sections were examined by confocal microscopy (Leica TCS SP5; Leica Microsystems), and digital images were taken, processed using the Leica Application suite program, and imported into a graphics program (Adobe Illustrator,) for display.

Expression of the canine *NPHP5* therapeutic transgene in the retina was visualized by RNA-*in situ* hybridization (RNA-ISH) RNAscope assay (Advanced Cell Diagnostics BioTechne). RNAscope probes against canine *NPHP5* (accession number: NM_001287550.1) were designed by ACDBio (catalog no. 462848), targeting the nucleotide regions 515-1592. As for IHC, ten micron thick sections from both treated and untreated regions of retinas were used, and target retrieval was performed on the sections by heating the slides in the Target Retrieval buffer at $88^0$C for 10 minutes followed by protease digestion. RNA-ISH was performed using the RNAscope 2.5 HD Assay-Red following the product manual guidelines. After hematoxylin staining, the sections were examined with a 40X objective and brightfield microscopy, and images captured as described above. Consecutive retinal sections were used for IHC, and capturing GFP fluorescence from the same retinal locations with epifluorescence microscopy (Axioplan, Carl Zeiss Meditec). Images digitally captured (Spot 4.0 camera; Diagnostic Instruments) and imported into Adobe Illustrator for display.

**Supplemental Figures**

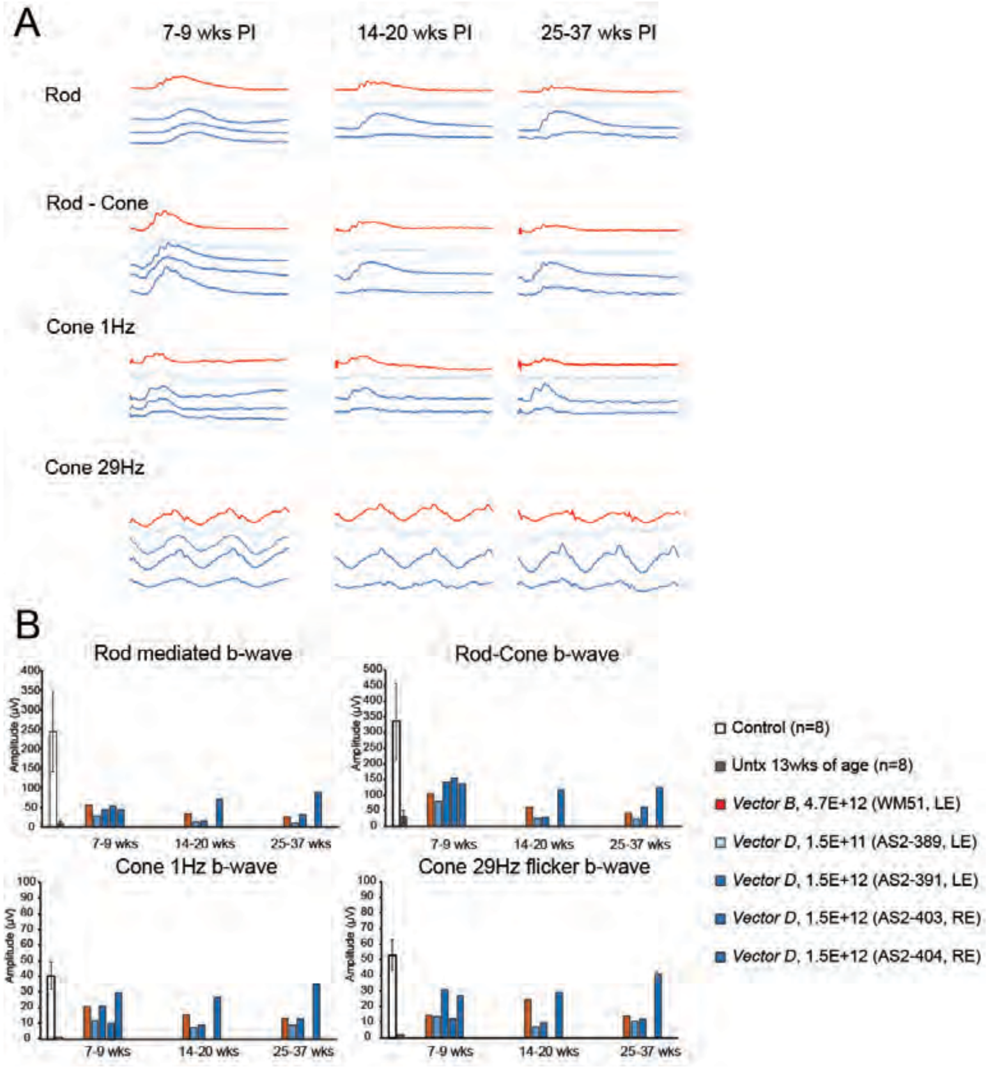

**Fig. S1. Robust and stable recovery of ERG rod and cone function following *NPHP5* gene augmentation therapy.** (**A**) ERG responses from the balance of the study dogs show the individual responses of five dogs treated with *Vectors B* and *D* at 3 different post injection (PI) intervals. All dogs show recovery of rod and cone ERG responses that remains stable for 6 months. (**B**) Summary data of the amplitudes recorded for individual treated dogs shown in (A) in response to rod, rod-cone and cone selective stimuli at 3 different PI intervals. Data for control (n=4 dogs/8 eyes; 22.2±3.4 wks of age) and untreated mutants (n=8; 13-14 wks of age) is provided for comparison and represents the mean± 1 SD of the amplitudes; the empty space in all graphs at the 14-20 and 25-32 wks time points is for dog AS2-403 that was removed for histopathology and IHC at 8 wks of PI. Light intensities used to elicit the illustrated responses are: Rod= -1.7 log cd·s·m–2; Rod Cone= 0.5 log cd·s·m–2; Cone 1Hz= 0.5 log cd·s·m–2; Cone 29Hz= 0.25 log cd·s·m–2. Vector titers are in vg/mL, and eyes received 70 $\mu$L subretinal injections. Individual animal identifiers are in parenthesis; LE-left eye; RE-right eye.

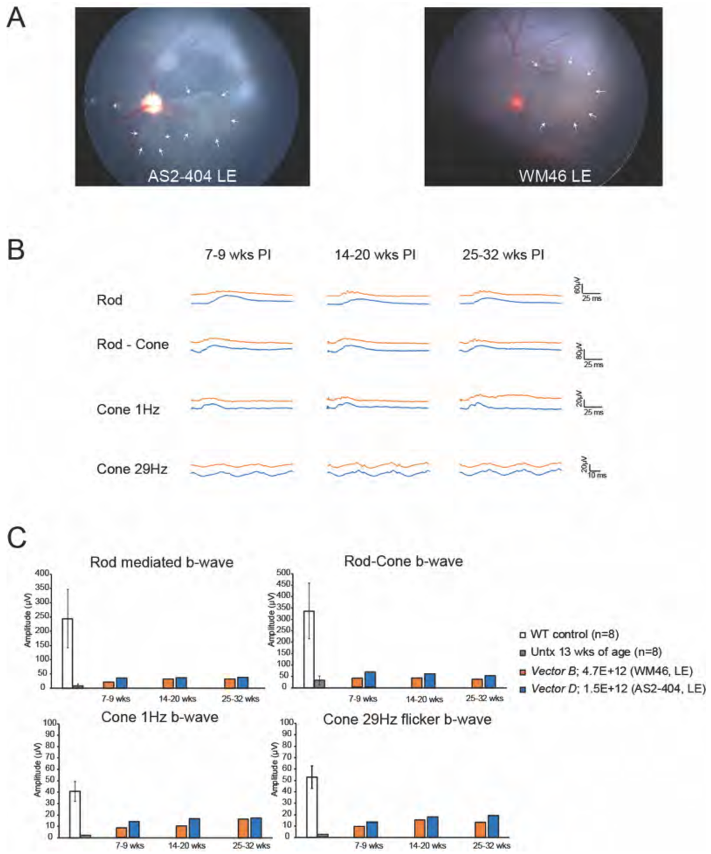

**Fig. S2. Stable functional recovery even with suboptimal treatments.** (**A**) RetCam fundus photographs taken of the subretinal blebs immediately after treatment in 2 eyes that had suboptimal blebs-multiloculated and flattened (left: AS2-404 LE) or with a large retinal tear (right: WM46 LE). (**B**) In spite of these surgical complications, there was recovery of rod and cone ERG function by 7-9 wks post injection (PI) which remained stable through the 25-32 wk study period. (**C**) The cone ERG amplitudes were comparatively higher than the rod responses. Data for control (n=4 dogs/8 eyes; 22.2±3.4 wks of age) and untreated mutants (n=8; 13-14 wks of age) is provided for comparison and represents the mean± 1 SD of the amplitudes Light intensities used to elicit the illustrated responses are: Rod= -1.7 log cd·s·m–2; Rod Cone= 0.5 log cd·s·m–2; Cone 1Hz= 0.5 log cd·s·m–2; Cone 29Hz= 0.25 log cd·s·m–2. Vector titers are in vg/mL, and eyes received 70 $\mu$L subretinal injections. Individual animal identifiers are in parenthesis; LE-left eye.

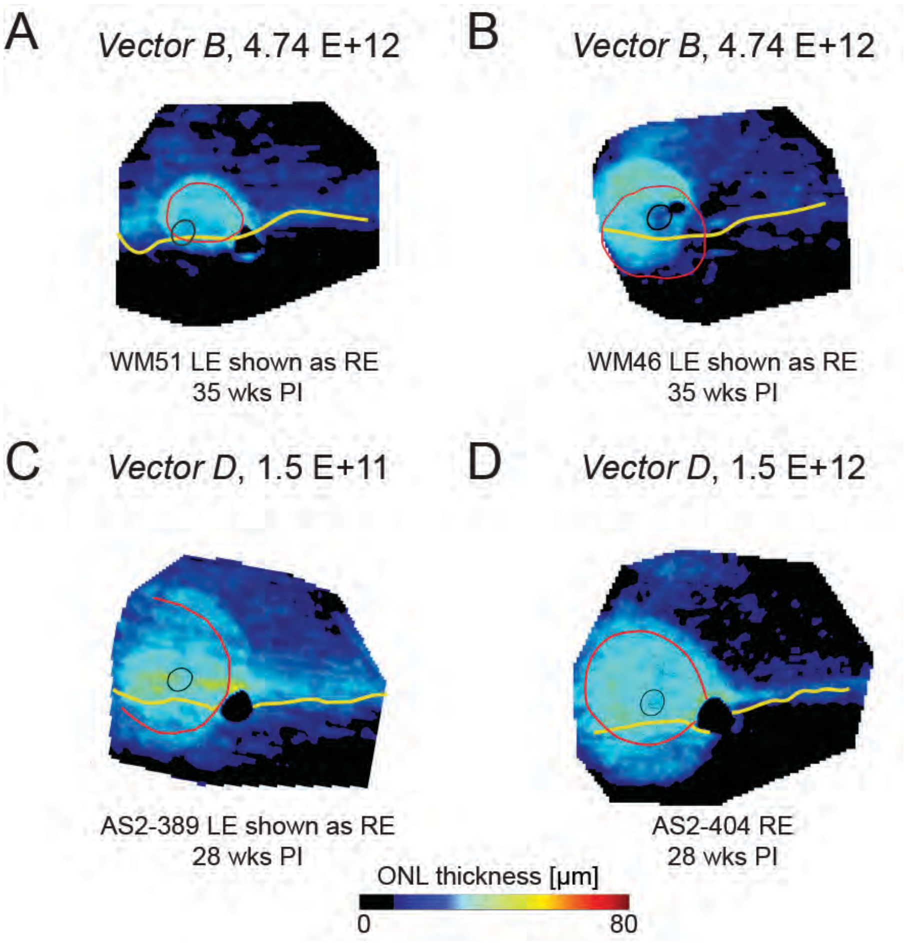

**Fig. S3. In vivo retinal imaging shows preservation of photoreceptor outer nuclear layer (ONL) in treated areas.** (**A** to **D**) Topographical maps of ONL thickness of the balance of study dogs. hapesAll eyes treated with the therapeutic vectors show preservation of ONL thickness in comparison to untreated areas of the same eye. Retained ONL thickness in some cases extends beyond the bleb border. Anatomic landmarks added include the fovea-like region (temporal black circle), the optic disc (black), and the tapetal boundary (wavy yellow line). In treated eyes, the red line identifies the subretinal injection bleb determined photographically immediately after the injection. LE-left eye; RE-right eye; PI-post injection interval. Vector titers are in vg/mL, and eyes received 70 $\mu$L subretinal injections.

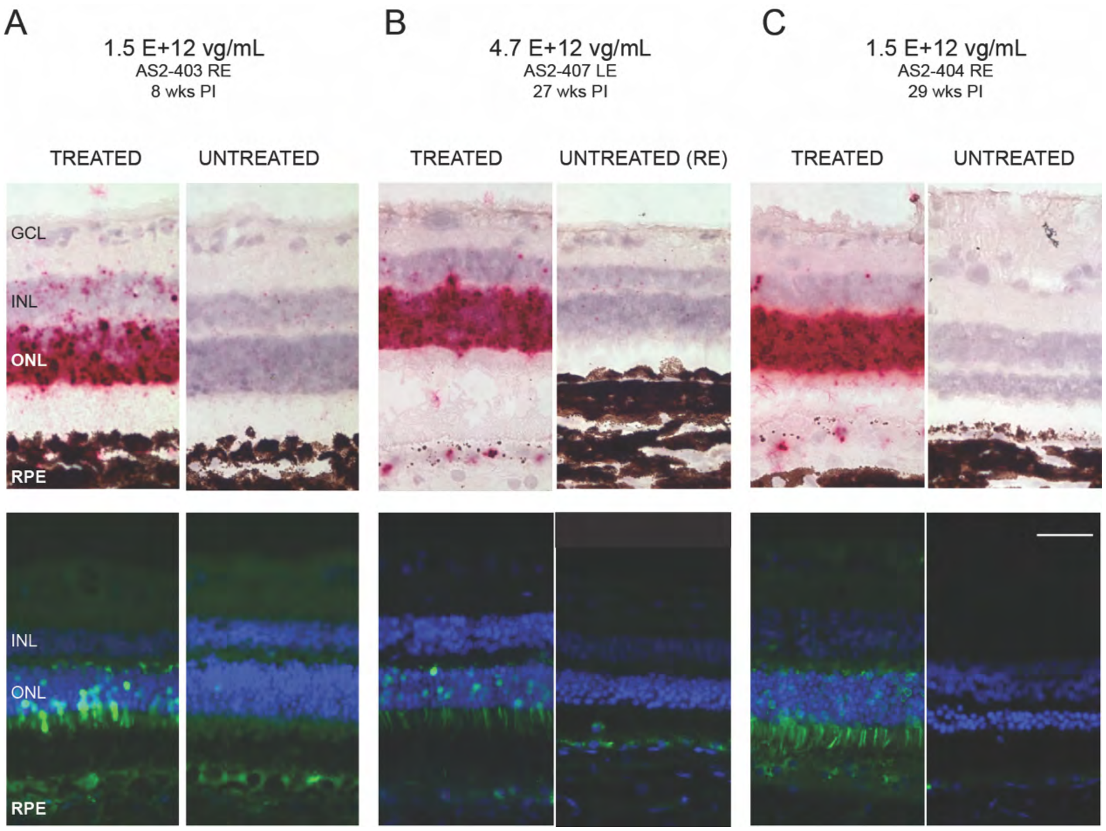

**Figure S4. Therapeutic *NPHP5* transgene is expressed only in treated area of injected eyes.** (**A** to **C**) In situ hybridization using an *NPHP5* RNA probe (top row) and GFP fluorescence from sequential sections taken from eyes subretinally-injected with the therapeutic *Vector D* 'spiked' with a GFP vector (table S1). Strong hybridization signals are present in treated areas, particularly the ONL at the earlier (A) and later (B, C) post injection (PI) time points. GFP fluorescence is observed in the photoreceptor inner segments and ONL of the treated areas. (*A-C*) Note that the ONL and photoreceptor layers of untreated regions thins at the later PI time points because of ongoing degeneration. Calibration marker in lower right figure=40 $\mu$m; GCL-ganglion cell layer, INL-inner nuclear layer, ONL-outer nuclear layer, RPE-retinal pigment epithelium, RE-right eye, LE-left eye, vg/mL=vector genomes/mL.

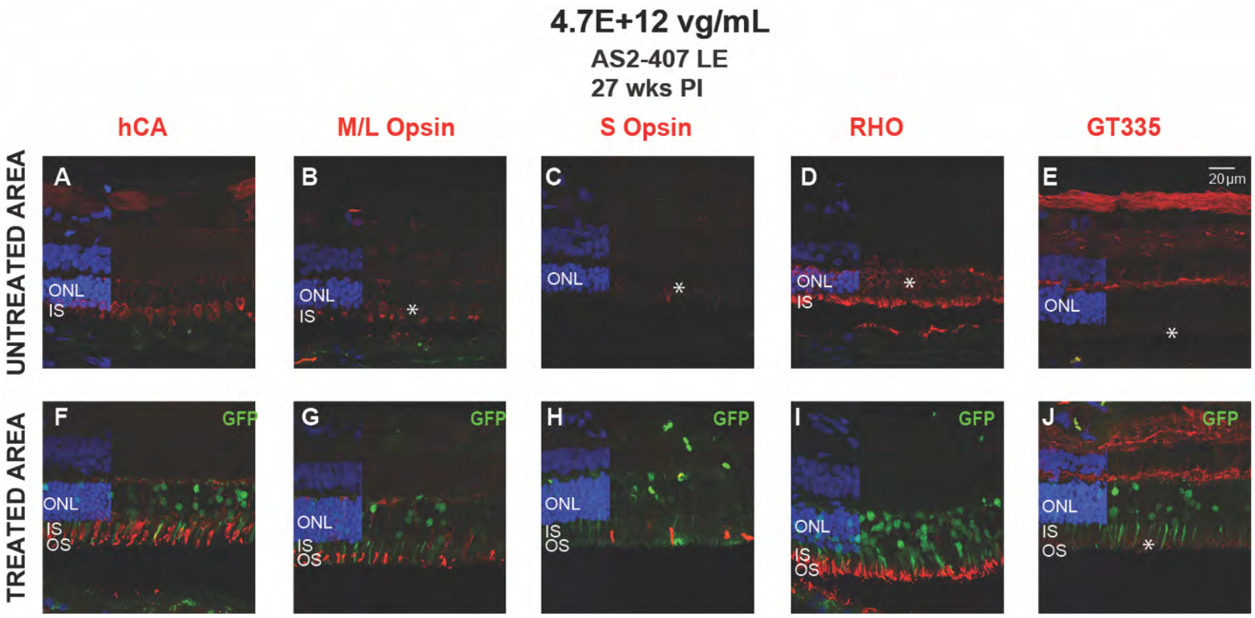

**Fig. S5. Stable recovery of normal rod and cone structure after treatment.** (**A** to **E**) Sections from untreated and (**F** to **J**) treated areas (*Vector D*; scAAV2/8(Y733F) GRK1-*cNPHP5*) of the left eye of dog AS2-407 at 27 weeks following therapy; treated areas identified by GFP fluorescence (green) in photoreceptor and outer nuclear layer nuclei. (A to C and F to H) Cone specific molecular markers show normal cone structure in treated areas, and all cone IS have corresponding OS. Untreated areas show absence of cone IS and OS, and cone opsin mislocalization to ONL (B and C*). (D and I) Rod OS are structurally normal, and opsin mislocalization is reversed in treated area, but in untreated area rod degeneration has advanced and remaining rod nuclei have perinuclear RHO localization (D*). (E and J) Localization of glutamylated *RPGR*$^{ORF15}$ in the ciliary transition zone (*) is distinct, while the untreated region has none. Hoechst nuclear label is used in all sections. ONL-outer nuclear layer; IS-inner segment; OS-outer segment; LE-left eye; vg/mL-vector genomes/mL.

**Movie S1: Restoration of Visual Function in Canine NPHP5 LCA by Gene Augmentation Therapy.** Functional vision evaluated at 28 weeks post injection in a dog treated at 5.1 weeks of age with *Vector C* (scAAV2/5-GRK1-hNPHP5) having the therapeutic human cDNA.

**table S1. Details of *NPHP5*-mutant dogs and vectors used in the study, and in vivo treatment outcomes.**

| Animal/Eye (Gender) | Age (wk) | | | Vector | | | Study week-PI interval (age if untreated) | | | *In vivo* Outcomes | |
|---|---|---|---|---|---|---|---|---|---|---|---|
| | Begin | End | PI interval | AAV construct (or BSS) | Vol (µL) | Titer (vg/mL) | ERG | OCT | Morph, IHC, ISH | ONL Rescue** (OCT) | ERG Rescue |
| AS-278 | - | 6 | - | - | - | - | 6 | - | 6 | – | – |
| AS2-387 | - | 15 | - | - | - | - | - | (15) | - | – | – |
| AS2-394 | | | | | | | | (14, 34) | | – | – |
| AS21-7/LE (F) | 6 | 33 | 28 | ssAAV2/5 IRBP-*cNPHP5** | 70 | LE:1.5E+12 | 7<br>14<br>25 | 8<br><br>27 | – | yes | yes |
| WM51/LE (F) | 5 | 41 | 36 | scAAV2/5-GRK1-*cNPHP5*<br><br>BSS | 70(20)<br><br>70 | LE: 4.7E+12<br>subRet (IVit)<br>RE: -- | 9<br>20<br>33 | <br>20<br>36 | – | yes | yes |
| WM46/BE (F) | 6 | 46 | 40 | scAAV2/5-GRK1-*cNPHP5* | 70(70)<br><br>70(30) | LE: 4.7E+12<br>subRet (IVit)<br>RE: 4.7E+12<br>subRet (IVit) | -0.3<br>8<br>19<br>37 | <br><br>21<br>40 | – | yes<br><br>yes | yes<br><br>yes |
| WM60/LE (F) | 5 | 35 | 30 | scAAV2/5-GRK1-***h****NPHP5*<br><br>BSS | 70<br><br>70 | LE: 4.7E+12<br><br>RE | 8<br>15<br>25 | <br>13<br>30 | – | yes | yes |
| AS2-396/BE (F) | 5 | 34 | 29 | scAAV2/8(Y733F) IRBP-*cNPHP5** | 70<br>70 | LE:1.5E+12<br>RE:1.5E+11 | 8<br>28 | 9<br>29 | 33 | yes<br>no | no<br>no |
| AS2-389/LE (F) | 6 | 33 | 28 | scAAV2/8(Y733F) GRK1-*cNPHP5** | 70 | LE:1.5E+11 | 7<br>14<br>25 | 8<br><br>27 | – | yes | yes |
| AS2-391/LE (F) | 6 | 33 | 28 | scAAV2/8(Y733F) GRK1-*cNPHP5** | 70 | LE:1.5E+12 | 7<br>14<br>25 | 8<br><br>27 | - | yes | yes |
| AS2-403/BE (M) | 6 | 14 | 8 | scAAV2/8(Y733F) GRK1-*cNPHP5** | 70 | BE:1.5E+12 | 7 | 8 | 8 | □ | yes-BE |
| AS2-404/BE (M) | 6 | 34 | 29 | scAAV2/8(Y733F) GRK1-*cNPHP5** | 70<br>70 | RE: 1.5E+12<br>LE: 4.7E+12 | 7<br>15<br>26 | 8<br><br>28 | 29 | yes<br>yes | yes<br>yes |
| AS2-407/LE (F) | 6 | 338 | 27 | scAAV2/8(Y733F) GRK1-*cNPHP5** | 70 | LE: 4.7E+12 | 7<br>15<br>26 | 8<br><br>27 | 27 | yes | yes |
| AS2-408 | - | 14 | - | - | - | - | BE: 14 | - | - | | |
| AS2-410 | - | 14 | - | - | - | - | BE: 14 | - | - | | |

* spiked with AAV2/5-IRBP-*GFP* at a titer of 5xE+10 vg/mL
** ONL rescue illustrated in Figure 4, or assessed from OCT volumetric scans
φ=PI too short to assess rescue

Vector designation in main text and in Fig. 2:
ssAAV2/5 IRBP-*cNPHP5* (*Vector A*)*;* scAAV2/5-GRK1-*cNPHP5* (*Vector B*)*;* scAAV2/5-GRK1-***h****NPHP5* (*Vector C*)*;* scAAV2/8(Y733F) GRK1-*cNPHP5* (*Vector D*)

Key:
cNPHP5 = canine *NPHP5*
ERG=electroretinography
GRK1=proximal human GRK1 promoter
hNPHP5 = human *NPHP5*
IHC=immunohistochemistry
IRBP = Interphotoreceptor Retinaldehyde Binding Protein
ISH=in situ hybridization
Morph=morphology
OCT=optical coherence tomography
BE=both eyes; RE=right eye; LE=left eye
PI=post injection interval
Vol=volume injected in µL
WT=wild type
– = not applicable

**table S2. Primary antibodies used in this study.**

| Antigen | Host | Source; Catalog No., or Name | Working Concentration | Specificity |
|---|---|---|---|---|
| Cone arrestin (human) | Goat polyclonal | Custom made; (Beltran Lab) | 1:50 | Cone photoreceptors |
| M/L opsin (human) | Rabbit polyclonal | Sigma-Aldrich, Chemicon®; AB5405 | 1:200 | M/L cones |
| S opsin (human) | Rabbit polyclonal | Sigma-Aldrich, Chemicon®; AB5407 | 1:200 | S cones |
| Rod opsin (rat) | Mouse monoclonal IgG1 | Sigma-Aldrich, Chemicon®; MAB5316 | 1:200 | rods |
| Glutamylated substrates | Mouse monoclonal IgG1 | AdipoGen Life Sciences; GT335 | 1:500 | Glutamylated RPGR in connecting cilium |
| eGFP | Mouse monoclonal IgG1 | Sigma-Aldrich, Chemicon®, MAB3580 | 1:1,000 | GFP reporter protein |
| **NPHP5 antibodies that showed no specific labeling against canine NPHP5** | | | | |
| NPHP5 (human) | Rabbit polyclonal | Santa Cruz Biotechnology; sc-134804 | 1:100-1:500 | Human NPHP5 residues 1-202 aa |
| NPHP5 (human) | Rabbit polyclonal | Novus Biologicals; NBP2-14126 | 1:100-1:500 | Human NPHP5 residues 420-480 aa |
| NPHP5 (human) | Rabbit polyclonal | IRCM1; (William Tsang Lab) | 1:100-1:500 | Human NPHP5 residues 469-598aa |
| NPHP5 (human) | Rabbit polyclonal | IRCM2, (William Tsang Lab) | 1:100-1:500 | Human NPHP5 residues 1-131aa |
